# Beta-cell-derived urocortin-3 contributes to glucose-regulated somatostatin and glucagon secretion in the mouse pancreas

**DOI:** 10.64898/2026.11.18.767593

**Authors:** Jon Vergara-Ucin, Daniel Bjørklund Andersen, Gianfranco Di Giuseppe, Teresa Mezza, Mette Marie Rosenkilde, Jens Juul Holst

## Abstract

**Aims/Hypothesis:** Glucagon secretion is impaired in both type 1 and type 2 diabetes. The mechanisms behind this dysregulation are incompletely understood. Urocortin-3 (UCN-3), a beta-cell-derived peptide, stimulates somatostatin (SST) secretion and may thereby regulate glucagon secretion. We investigated the mechanisms through which UCN-3 regulates pancreatic hormone secretion and examined the physiological contribution of endogenous UCN-3 to glucose-regulated hormone secretion.

**Methods:** Pancreatic hormone secretion was studied using the isolated perfused mouse pancreas. The effects of exogenous and endogenous UCN-3 were investigated using selective antagonists of SST receptors (SSTR) and corticotropin-releasing hormone receptor 2 (CRHR2). In addition, the role of UCN-3 was examined in streptozotocin (STZ) treated mice.

**Results:** Exogenous UCN-3stimulated somatostatin secretion and inhibited glucagon secretion in the mouse pancreas. Blockade of SSTRs completely abolished the glucagonostatic effect of UCN-3, demonstrating that inhibition of glucagon secretion is indirectly mediated through SST signaling. Moreover, CRHR2 antagonism prevented both UCN-3-induced SST secretion and inhibition of glucagon secretion, identifying CRHR2 as the key receptor mediating these effects. During acute glucose stimulation (3.5-10mM), CRHR2 antagonism reduced glucose-induced stimulation of SST and attenuated glucagon inhibition, indicating that endogenous UCN-3 participates in the acute islet response to hyperglycemia. However, during sustained hyperglycemia, CRHR2 antagonism reduced SST secretion without altering glucagon secretion. In STZ-treated mice, glucose-induced SST secretion was lost.

**Conclusions:** Urocortin-3 regulated glucagon secretion indirectly through CRHR2-dependent stimulation of SST secretion. Endogenous UCN-3 contributes to the acute islet response to hyperglycemia, supporting a physiologically relevant beta-cell-to-delta-cell-to-alpha-cell paracrine axis in the mouse pancreas. This pathway is disrupted following beta-cell loss, suggesting a potential contribution to impaired islet hormone regulation in diabetes.

## Introduction

Glucagon, secreted from pancreatic alpha-cells, plays an essential role in glucose homeostasis by promoting hepatic glucose production during fasting [1]. Appropriate regulation of glucagon secretion is therefore critical for maintaining normal blood glucose levels. Dysregulated glucagon secretion is a characteristic feature of both type 1 diabetes and type 2 diabetes. Patients with type 1 diabetes lose the ability to appropriately increase glucagon secretion during hypoglycemia [2] whereas patients with type 2 diabetes often develop hyperglucagonemia, resulting in excessive hepatic glucose production, which contributes to hyperglycemia [3]. Although these abnormalities have been recognized for decades, the mechanisms responsible for the dysregulation of alpha-cell function remain incompletely understood.

Alpha-cell activity is regulated not only by nutrients but also by a complex network of intra-islet paracrine signals [4, 5]. Among these, somatostatin (SST), secreted from pancreatic delta-cells, is considered one of the most important inhibitory regulators of glucagon secretion. Alpha-cells express high levels of somatostatin receptor 2 (SSTR2), a Gαi-coupled receptor that suppresses cAMP signaling and potently inhibits glucagon secretion [6, 7]. Consequently, delta-cells are increasingly recognized as key integrators of intra-islet signaling networks, coordinating both insulin and glucagon secretion through SST release[8]. Several beta-cell-derived factors have also been proposed to influence alpha-cell activity, including insulin [9, 10], zinc [11], GABA [12], and other co-secreted peptides. While some studies have suggested that insulin directly or indirectly inhibits glucagon secretion, the physiological importance of this mechanism remains debated [13].

One beta-cell-derived peptide that has attracted considerable interest is urocortin-3 (UCN-3). UCN-3 is among the most abundant peptides expressed by mature beta-cells and is co-secreted with insulin in response to glucose stimulation [14]. Previous studies have identified the corticotropin-releasing hormone receptor 2 (CRHR2) as the cognate receptor for UCN-3 in the endocrine pancreas. CRHR2 expression has been reported to be restricted to pancreatic delta-cells, where receptor activation stimulates Gαs signaling, increases intracellular cAMP, and promotes SST secretion [14, 15]. Through this mechanism, UCN-3 has been shown to participate in an inhibitory feedback loop to modulate insulin secretion in response to high glucose.

There is agreement that UCN-3 may influence delta-cell activity; however, the contribution to glucagon secretion remains less well understood. Previous studies have demonstrated that UCN-3 decreases Ca^2+^ and cAMP in alpha-cells and inhibits glucagon secretion in mouse islets [16] and rat pancreas [17]. However, while the pharmacological actions of exogenous UCN-3 have been characterized, it remains unclear whether endogenous UCN-3 contributes to the physiological regulation of pancreatic hormone secretion during glycemic changes. Therefore, in the present study, we used the isolated perfused mouse pancreas to investigate the mechanisms by which UCN-3 regulates glucagon and SST secretion and to determine the physiological contribution of endogenous UCN-3 signaling to these secretions during glycemic changes.

## Materials and methods

### Animals

The experiments were conducted in accordance with the guidelines of Danish legislation governing animal experimentation (1987) and with permission from the Danish Animal Experiments Inspectorate (2023-15-0201-01409). The animals were 10-15-week-old C57BL/6J male mice purchased from Janvier Lab. Mice were housed 8 per cage with a 12-hour light-dark cycle and free access to a chow diet and water.

To generate a beta-cell-deficient mouse model, 10-week-old C57BL/6J mice were treated with a single injection of streptozotocin (200mg/kg, intraperitoneal (i.p)) (Sigma-Aldrich, catalog number S0130) dissolved in sodium citrate buffer (pH 4.5) one week before the experimental day. Water supplemented with 10% sucrose was provided to the animals for 48 hours after the injection to prevent hypoglycemic events, and body weight was measured daily to assess disease severity. On the day of the experiment, blood glucose was measured to confirm diabetic phenotype. No animals reached predefined humane endpoints prior to completion of the experiment.

### Isolation and perfusion of the mouse pancreas

The perfusion method is described elsewhere [18]. In short, mice were anesthetized with an i.p. injection of ketamine/xylazine ((0.1 mL/20g), ketamine (Ketaminol®, Intervet, Merck & CO, cat. no. 511485), and Xylazine (Rompun® vet, Bayer AG, cat. no. 148999)) and placed on the surgical table heated at 37°C. An incision was made to open the abdominal cavity. Then, the small and large intestines (except for the most proximal part of the duodenum), the stomach, and the spleen were removed, with all supplying vasculature ligated. The renal blood supply was also ligated. A catheter was then inserted in the abdominal aorta, and perfusion of the organ was started. The aorta was ligated in the thoracic region to avoid perfusion of the heart, and the collecting catheter was placed in the portal vein from which the effluent was collected. The pancreas was perfused at a flow rate of 1.5 or 0.75 mL/min, and samples were collected every minute or every other minute, respectively. A 30-minute wash period was included before the start of the experiment. Samples were collected, placed immediately on ice, and stored at -20°C until further analysis.

### Solutions and test compounds

The solution used to perfuse the pancreases during the experiments was a modified Krebs-Ringer buffer (118.4mM NaCl, 2.95mM KCl, 2.58mM CaCl_2_, 1.22mM MgSO_4_, 1.18mM KH_2_PO_4_, 25mM NaHCO_3_) (supplemented with 0.1% Bovine Serum Albumin (BSA) (Merck, cat no. 112018), 5% Dextran-70 (Pharmacosmos, cat. no. 5510 0070 9001), 5 mM pyruvate (ThermoScientific, cat no. J61840), and 3 mM Vamin® (Fresenius-Kabi, cat. no.3017141)) that was gassed with 95% O2 and 5% CO2.

UCN-3, H6056 (SSTR2a), and H-5884 (SSTR5a) were purchased from Bachem (catalog numbers 4040194, 4044525, and 4041641). CYN 154806 (SSTR2a) was ordered from Sigma-Aldrich (catalog number C2490), and MK-4256 (SSTR3a) was purchased from MedChem Express (catalog number HY-13466). K41498 (CRHR2 antagonist) was purchased from Tocris Biosciences (catalog number 2070/1). These compounds were diluted in Milli-Q + 1% BSA and stored at -20°C. On the day of the experiment, the stock solutions were diluted in perfusion buffer to the desired concentration. L-Arginine (L-Arg) was purchased from Sigma-Aldrich (Germany; catalog number A6969), dissolved in perfusion buffer, and aliquots were stored in the freezer until the experimental day.

### Cell transfection

Cell transfection was carried out as previously described [19]. In short, 1.5 million HEK293 cells were seeded into T25 flasks in a growth medium consisting of DMEM + 10% Fetal Bovine Serum (FBS) and 1% Penicillin-Streptomycin. The following day, cells were transfected using the calcium phosphate transfection method, with 10 µg of plasmid containing mouse CRHR2 (mCRHR2) DNA or an empty vector. On the third day, the cell medium was removed from the flask, and the cells were washed with PBS. Then, cells were incubated for 2 minutes with 2mL Versene solution and supplemented with 3mL cell medium. Everything was mixed gently using a serological pipette, transferred to a 15mL tube, and centrifuged for 5 min at 1500rpm. The supernatant was removed, and cells were resuspended in 2mL of cell culture medium before immunostaining.

### Immunostaining

Concentrations and information of all antibodies used are provided in Supplemental Table 1. Mice were euthanized by cervical dislocation; the pancreas was dissected and fixed with 4% paraformaldehyde (PFA) overnight, dehydrated in 70% ethanol (EtOH), and embedded in paraffin. All paraffin-embedded tissue received the same pretreatment, and both pretreatment and staining were carried out on the same days. First, sections were deparaffinized by heating for 1 hour at 60°C, then immersed in Histolab-Clear (Histolab Products AB, Cat. No. 14250), followed by two incubations in separate HistoClear baths for 8min each. Then sections were rehydrated in a series of ethanol baths of decreasing concentrations (99%, 99%, 99%, 96%, 96%, 70%). All sections were then left in running water for 5 minutes. Following this, sections were placed in boiling citrate buffer (pH 6) and allowed to cool for 1 hour, or until the buffer reached 40°C. Then, sections were blocked with 2% BSA for 10 minutes, and primary antibodies were added at the desired concentrations and incubated at 4°C overnight. On the second day, all sections were washed 3 times with PBS, incubated with the secondary antibody, coverslips were mounted using Fluoroshield mounting medium with DAPI (Cat. No. ab104139, Abcam, Cambridge, UK), and sections were stored at 4°C. For CRHR2 staining, the signal was amplified using a VectaFluor amplification kit (Anti-rabbit, 488, Cat. no.: DK-1488, Vector Laboratories, California, USA), following the supplier’s instructions.

HEK293 cells transfected with either mCRHR2 DNA or an empty vector were seeded into a 24-well plate coated with poly-D-lysine and incubated overnight at 37°C in cell culture medium. Then, the medium was aspirated, and cells were washed twice with PBS, followed by fixation with 4% PFA for 10 minutes. Cells were washed again and blocked for 10 minutes in 2% BSA. Then, cells were incubated with primary anti-CRHR2 antibody overnight at 4°C, followed by staining with the same VectaFluor amplification kit used for the pancreas sections (Anti-rabbit, 488, Cat. no.: DK-1488, Vector Laboratories, California, USA), according to the supplier’s instructions.

### Image analysis and quantification

Fluorescence images were acquired on a Zeiss Axioscan 7 slide scanner using identical acquisition settings for all the slides. Images were analyzed in QuPath (version 0.5.1). Tissue area was defined by thresholding the DAPI channel, and insulin-positive area was defined by thresholding the Alexa Fluor 488 channel. The same fixed thresholds were applied to all images. Areas with debris, folds, or tears were manually excluded from the analysis. This analysis was performed blindly to minimize bias. The insulin-positive area was expressed as a ratio of the total tissue area and was normalized to the control group mean.

### Hormone measurements

Glucagon and somatostatin from the perfusate samples were measured using in-house radioimmunoassays (protocol and antibody code names: 4305 [20] and 1758 [21], respectively).

### Data presentation and statistical analysis

All data are presented as mean + SEM. Hormone output (fmol/min) was analyzed for each animal across the experimental periods. For each period, the area under the curve (AUC, fmol/10min) was calculated over the final 10 minutes of each period using the trapezoidal rule. Missing samples within a window were omitted, and AUC was calculated with the remaining points. Calculations were performed in R (version 3.5.2). Statistical significance was assessed using a paired T-test or a mixed-effects analysis followed by a Holm-Šídák correction for multiple comparisons. No animals or datasets were excluded other than perfusions lacking a positive control response to L-arginine. Because each pancreas served as its own control, randomization was not performed. Sample sizes were based on previous perfusion experiments from our group using similar experimental endpoints and were not determined by a power calculation. Perfusion experiments and hormone measurements were performed unblinded.

For image analysis, data are presented as mean ± SEM, with each data point representing one animal. Differences in insulin-positive area between control and STZ-treated mice were assessed using Welch’s T-test.

Statistical tests were performed using GraphPad Prism v10.6.1 (La Jolla, CA). Graphs were constructed in GraphPad Prism, and figures were assembled using Adobe Illustrator. P<0.05 was considered significant.

## Results

### Exogenous urocortin-3 stimulates somatostatin and inhibits glucagon secretion in the mouse pancreas

To first assess the effects of urocortin-3 on glucagon and SST secretion, we performed repeated infusions of the peptide (10nM) into the perfused mouse pancreas at low glucose (3.5mM) and measured perfusate hormone levels. UCN-3 strongly stimulated SST secretion during both infusions (1^st^ Infusion: basal 39.21±7.25 fmol/10min, vs UCN-3 171.32±33.63 fmol/10min, p-value < 0.05; 2^nd^ Infusion: basal 59.67±13.63 fmol/10min vs UCN-3 143.46±28.43fmol/10min, p-value<0.05) (Fig. 1A-B). Glucagon secretion, by contrast, was inhibited by UCN-3 (1^st^ infusion: basal 248.75±48.61 fmol/10min vs UCN-3 163.5±36.8 fmol/10min, p-value<0.05; 2^nd^ infusion: basal 210.75±33.73 fmol/10min vs UCN-3 134.87±44.4 fmol/10min, p-value<0.05) (Fig. 1C-D). These data confirm that the regulatory effects of UCN-3 previously described [16, 17] are also present in our experimental model.

**Figure 1.**
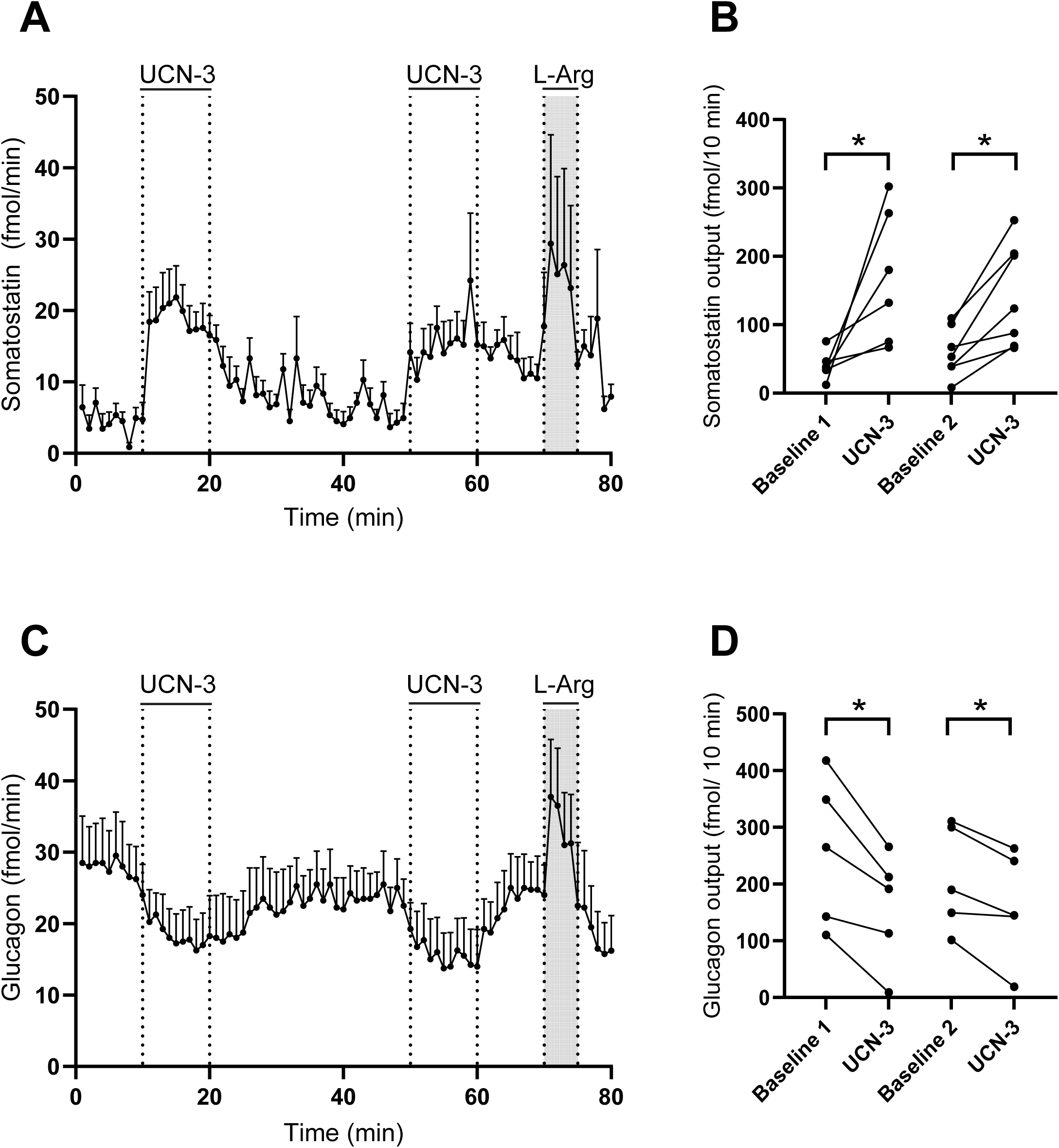
Urocortin-3 stimulates somatostatin and inhibits glucagon secretion in the perfused mouse pancreas. Somatostatin (A) and glucagon (C) secretion from isolated perfused mouse pancreas in response to 10nM UCN-3 infusion. SST (B) and glucagon (D) output. Baseline buffer has 3.5mM glucose. Data are presented as mean + SEM in A and C, and as individual data points in B and D. N(A, B)=6; N(C, D)=5. Statistical significance was assessed using a mixed-effects analysis, followed by a Holm-Šídák correction for multiple comparisons (*P<0.05).

### Exogenous urocortin-3 indirectly inhibits glucagon secretion through SST receptor activation

Once the effects of UCN-3 in the mouse pancreas were defined, we next tested whether the inhibitory effect on glucagon secretion was direct or indirect via stimulation of SST secretion. To test this, we again performed a double infusion of UCN-3 at 3.5 mM glucose, but this time co-infused with SST receptor (SSTR) antagonists (SSTRa) during the second infusion to block the most prevalent pancreatic SST receptors (SSTR2, 3, and 5).

Again, UCN-3 alone stimulated SST secretion (103.59±12.04 fmol/10min vs 316.68±56.16 fmol/10min, p-value<0.05) (Fig. 2A-B) and inhibited glucagon secretion (222.85±65.82 fmol/10min vs 148.6±53.79, p-value<0.05) (Fig. 2C-D). Blocking SST actions increased the secretion of both glucagon and somatostatin, demonstrating a tonic inhibitory action of SST over both alpha- and delta-cells even at low glucose levels. SSTR antagonism prevented UCN-3-induced glucagon inhibition (239.68±67.57 fmol/10min vs 300.32±102.05 fmol/10min, p-value=0.25) (Fig. 2C-D) and further stimulated SST secretion (326.81±48.79 fmol/10min vs 703.5±104.42 fmol/10min, p-value<0.001) (Fig. 2A-B). These findings demonstrate that UCN-3 indirectly inhibits glucagon secretion via SST receptor activation.

**Figure 2.**
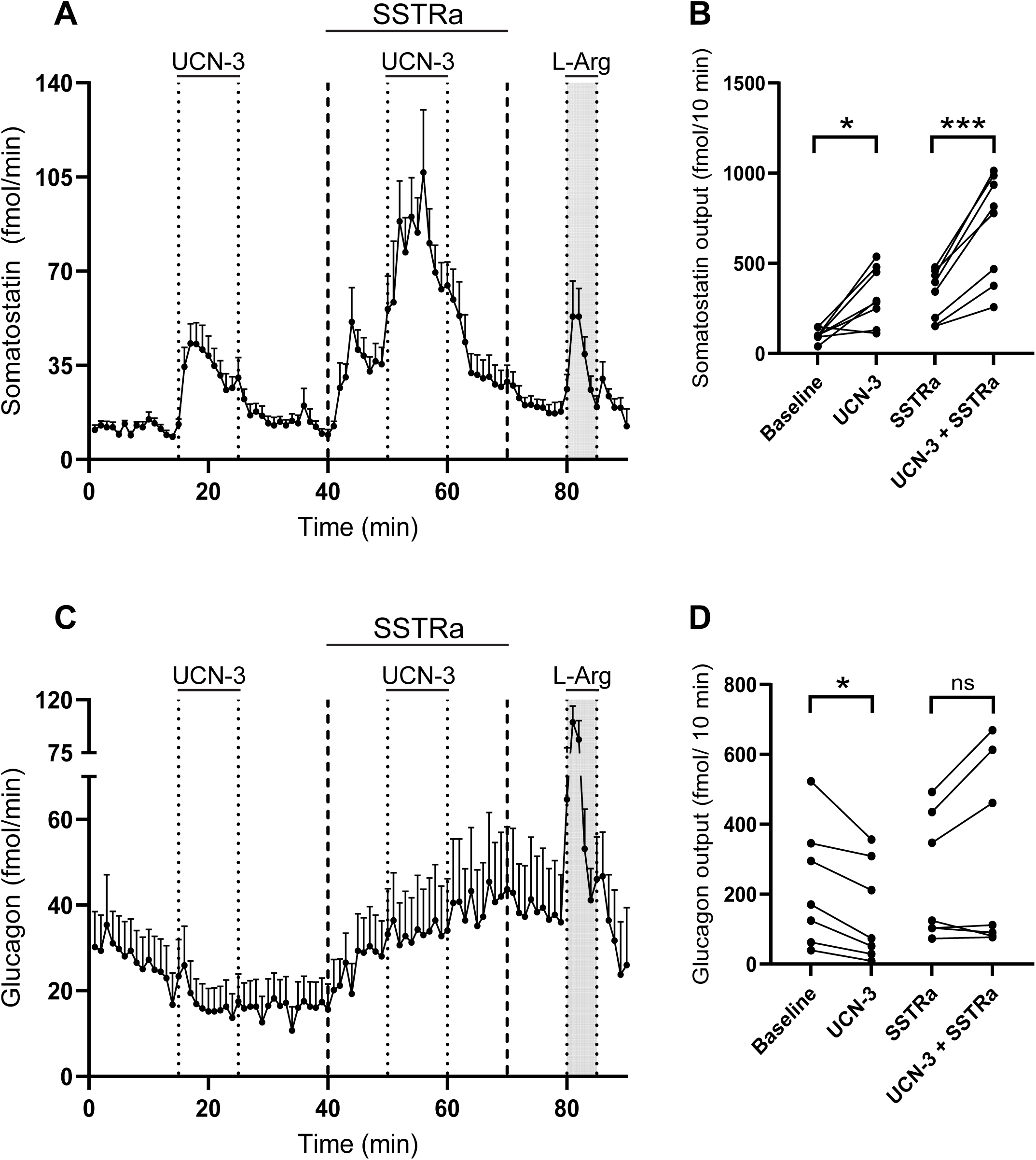
SST receptor activation is essential for UCN-3-induced inhibition of glucagon secretion. Somatostatin (A) and glucagon (C) secretion from the perfused mouse pancreas in response to 10nM UCN-3 with or without a cocktail of SSTR antagonists (100nM of each antagonist, 400nM in total). SST (B) and glucagon (D) output. Baseline buffer has 3.5mM glucose. Data are presented as mean + SEM in A and C, and as individual points in B and D. N=6. Statistical significance was assessed by mixed-effects analysis followed by Holm-Šídák correction for multiple comparisons (*P<0.05, ***P<0.001, ns – non-significant).

### CRHR2 activation is necessary for the somatostatin stimulation driven by exogenous UCN-3

CRHR2 is the primary receptor mediating UCN-3 signaling. Previous studies reported that CRHR2 is expressed exclusively in delta-cells in mouse pancreatic islets, as determined by qPCR or in situ hybridization. Here, to investigate CRHR2 localization, we co-immunostained mouse pancreatic sections with anti-SST and anti-CRHR2 antibodies. First, we tested the specificity of the anti-CRHR2 antibody by staining HEK293 cells transfected with either mouse CRHR2 (mCRHR2) or a vehicle DNA plasmid. Only the HEK293 cells transfected with mCRHR2 DNA showed a fluorescent signal (Suppl. Fig. 1), supporting the antibody’s ability to recognize recombinant mCRHR2. In mouse pancreatic sections, the CRHR2 immunoreactivity completely overlapped with SST-positive cells, consistent with reports showing selective Crhr2 transcript expression in pancreatic delta-cells (Fig. 3A).

**Figure 3.**
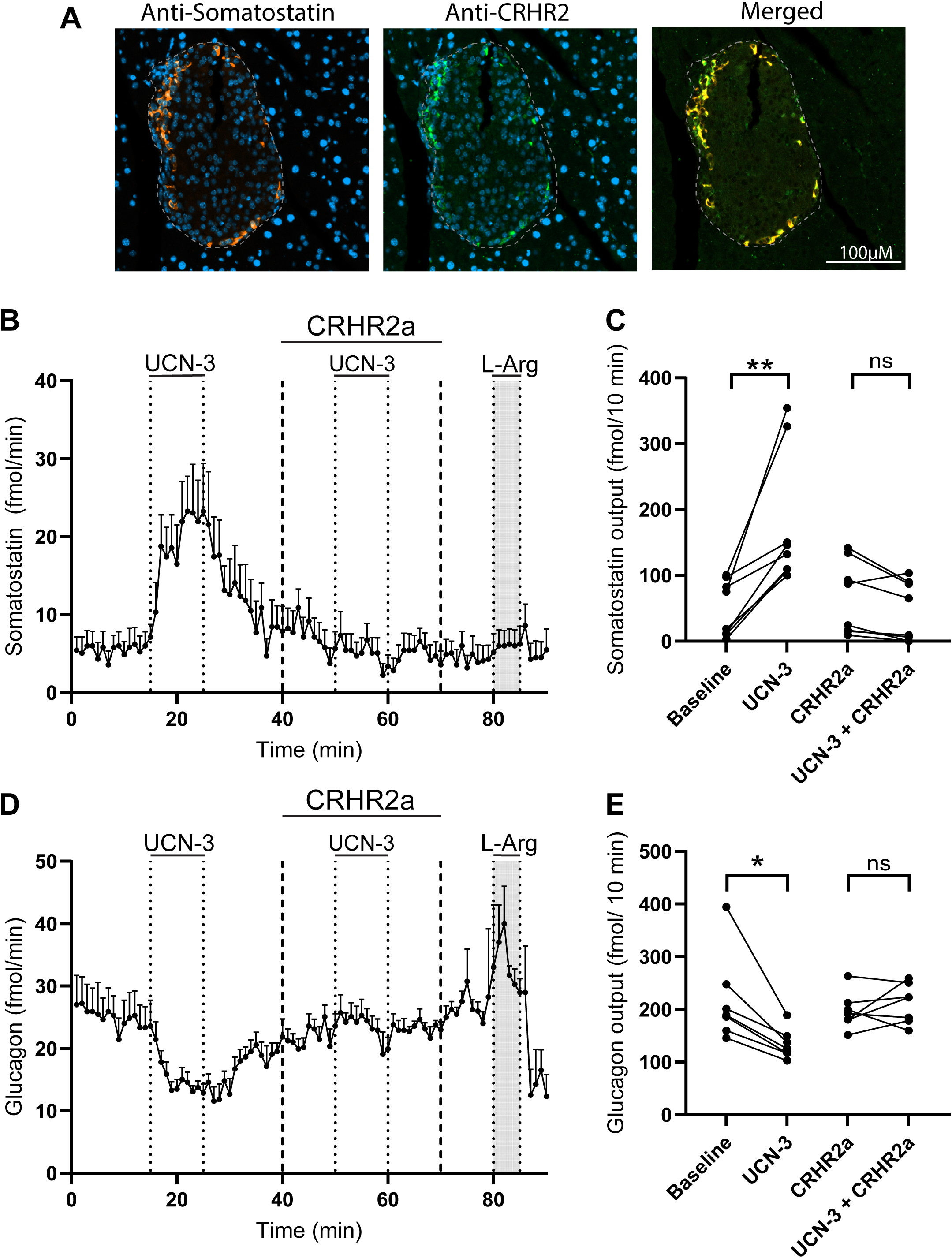
CRHR2 antagonism prevents UCN-3 actions in the endocrine pancreas. Representative mouse islet section co-immunostained for somatostatin and CRHR2, together with the merged image (A). SST (B) and glucagon (D) secretion from the perfusate in response to 10nM UCN-3 with or without the CRHR2 antagonist (K41498, 100nM). SST (C) and glucagon (E) output. Baseline buffer has 3.5mM glucose. Data are presented as mean + SEM in B and D, and as individual points in C and E. N=6. Statistical significance was assessed by mixed-effects analysis followed by Holm-Šídák correction for multiple comparisons (*P<0.05,**P<0.01, ns – non-significant).

Next, we studied whether CRHR2 is essential for UCN-3 to influence pancreatic hormone secretion, or whether UCN-3 acts through other mechanisms. We performed sequential infusions of UCN-3 in the absence or presence of a specific CRHR2 antagonist (CRHR2a). Once again, when infused alone, UCN-3 increased SST (49.96±15.04 fmol/10min vs 178.21±36.0 fmol/10min, p-value<0.01) (Fig. 3B-C) and lowered glucagon secretion (217.6±31.93 fmol/10min vs 133.93±10.8 fmol/10min, p-value<0.05) (Fig. 3D-E), but K41498 fully prevented the effects on SST (65.06±19.65 fmol/10min vs 45.09±16.08 fmol/10min, p-value=0.08) (Fig. 3B-C), and glucagon (197.03±13.16 fmol/10min vs 211.17±14.24 fmol/10min, p-value=0.61) (Fig. 3D-E). Together, our data support a mode of action in which exogenous UCN-3 specifically stimulates SST secretion through CRHR2, which in turn inhibits glucagon secretion through SSTR.

### Endogenous urocortin-3 participates in glucose-regulated pancreatic hormone secretion

Our previous experiments investigated the regulatory effects of exogenous UCN-3 on somatostatin and glucagon secretion at low glucose. However, *in vivo*, endogenous UCN-3 would be secreted when beta-cells are activated, which is most prominent at glucose concentrations above 6 mM, meaning that UCN-3 would not be actively secreted under the previously tested conditions. We therefore investigated whether endogenous UCN-3 is involved in the hyperglycemia-induced inhibition of glucagon through SST stimulation. This was tested by increasing glucose levels from 3.5 mM to 10 mM in the absence or presence of K41498. As expected, increasing glucose levels stimulated SST secretion (43.65±3.41 fmol/10min vs 74.4±8.45 fmol/10min, p-value<0.05) (Fig. 4A-B) and inhibited glucagon secretion (129.3±15.75 fmol/10min vs 70.65±7.94 fmol/10min, p-value<0.01) (Fig. 4C-D). In the presence of K41498, hyperglycemia still numerically reduced glucagon secretion (106.8±8.59 fmol/10min vs 92.85±6.75 fmol/10min, p-value=0.29) (Fig. 4C-D) and increased SST secretion (32.25±4.44 fmol/10min vs 38.85±5.74 fmol/10min, p-value=0.305) (Fig. 4A-B), although neither effect reached statistical significance; however, the effect size was reduced (Glucagon: 44.8% inhibition without vs 11.4% with CRHR2a, p-value<0.01; SST: 71% stimulation without vs 24% stimulation with CRHR2a, P-value=0.11), indicating that endogenous UCN-3 signaling is a physiologically relevant component of the acute islet response to hyperglycemia.

**Figure 4.**
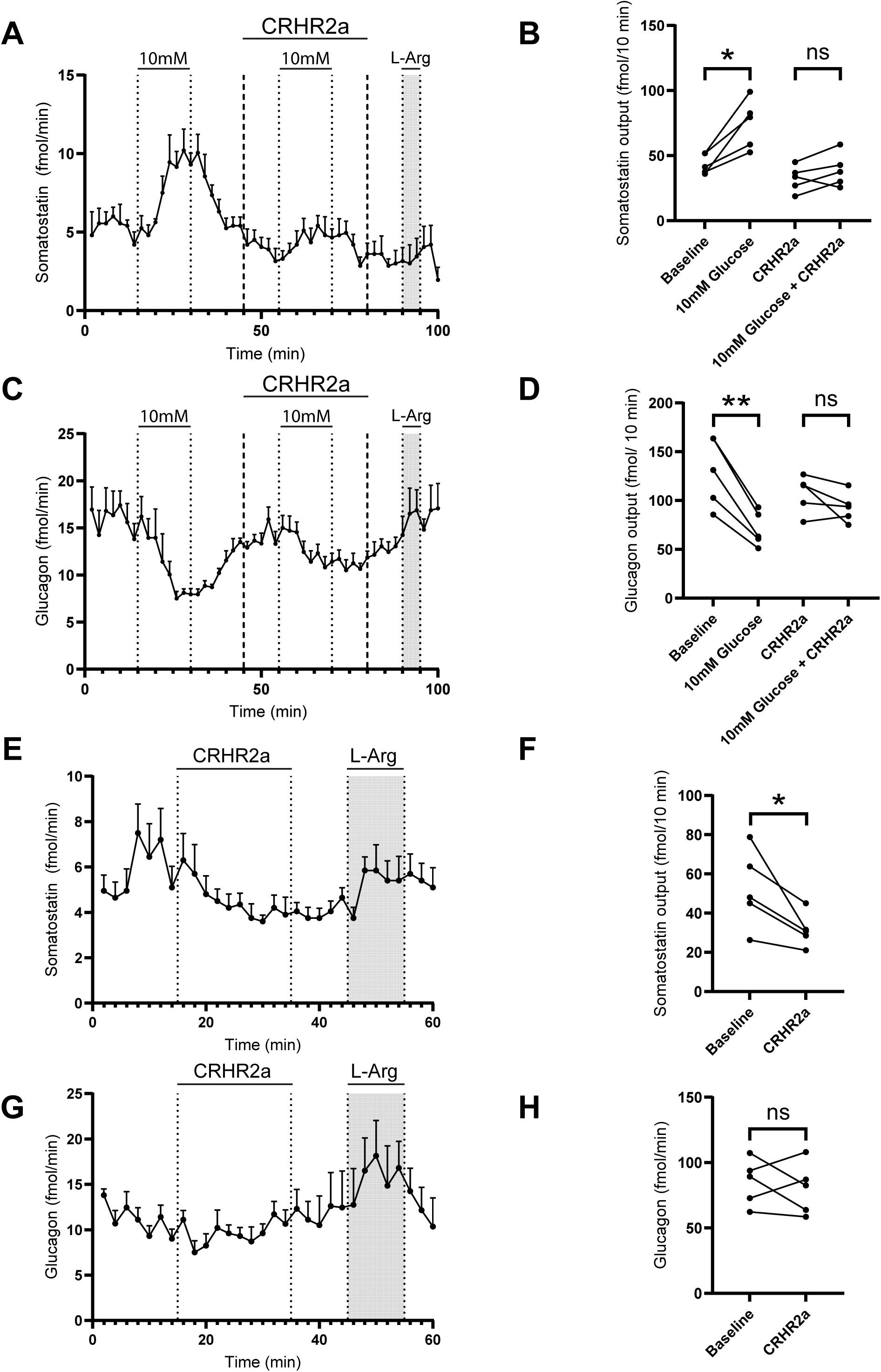
Endogenous urocortin-3 contributes to the acute response to hyperglycemia, whereas its role during sustained hyperglycemia is restricted to somatostatin secretion. Somatostatin. (A) and glucagon (C) secretion from the perfused mouse pancreas in response to high glucose (10mM) with or without the CRHR2 antagonist (K41498, 100nM). SST (B) and glucagon (D) output. For A and C, the baseline buffer is 3.5mM glucose. SST (E) and glucagon (G) secretion from the mouse-perfused pancreas at 10mM glucose levels in response to CRHR2 antagonist (K41498, 100nM). SST (F) and glucagon output (H). In E and G, the baseline buffer contains 10 mM glucose. Data are presented as mean + SEM for A, C, E, and G, and as individual points in B, D, F, and H. N=5. Statistical significance was assessed by mixed-effects analysis followed by Holm-Šídák correction for multiple comparisons in B and D, and by a paired T-test in F and H. (*P<0.05, **P<0.01, ns – non-significant).

However, when the pancreas was continuously perfused under hyperglycemic conditions (10 mM), blocking CRHR2 and therefore blocking endogenous UCN-3 signaling reduced SST secretion (48±8.46 fmol/10min vs 32.5±3.37 fmol/10min, p-value<0.05) (Fig. 4E-F), while it did not affect glucagon secretion (78.37±9.29 fmol/10min vs 76.5±8.00 fmol/10min, p-value=0.59) (Fig. 4G-H).

Together, our data indicate that endogenous UCN-3 signaling is a physiologically relevant component of the acute glucagon and SST response to hyperglycemia, but becomes less impactful in the setting of chronic hyperglycemia.

### Disruption of the beta-cell paracrine environment impairs glucose-regulated hormone secretion

Given this, we investigated this paracrine axis in a mouse model with reduced UCN-3 signaling, the streptozotocin (STZ)–treated mice. Intraperitoneal injection of a single high dose of STZ leads to beta-cell ablation (approximately 80% reduction, P<0.0001) (Fig. 5A), allowing us to investigate pancreatic hormone secretion in a model with a reduced beta-cell paracrine environment.

**Figure 5.**
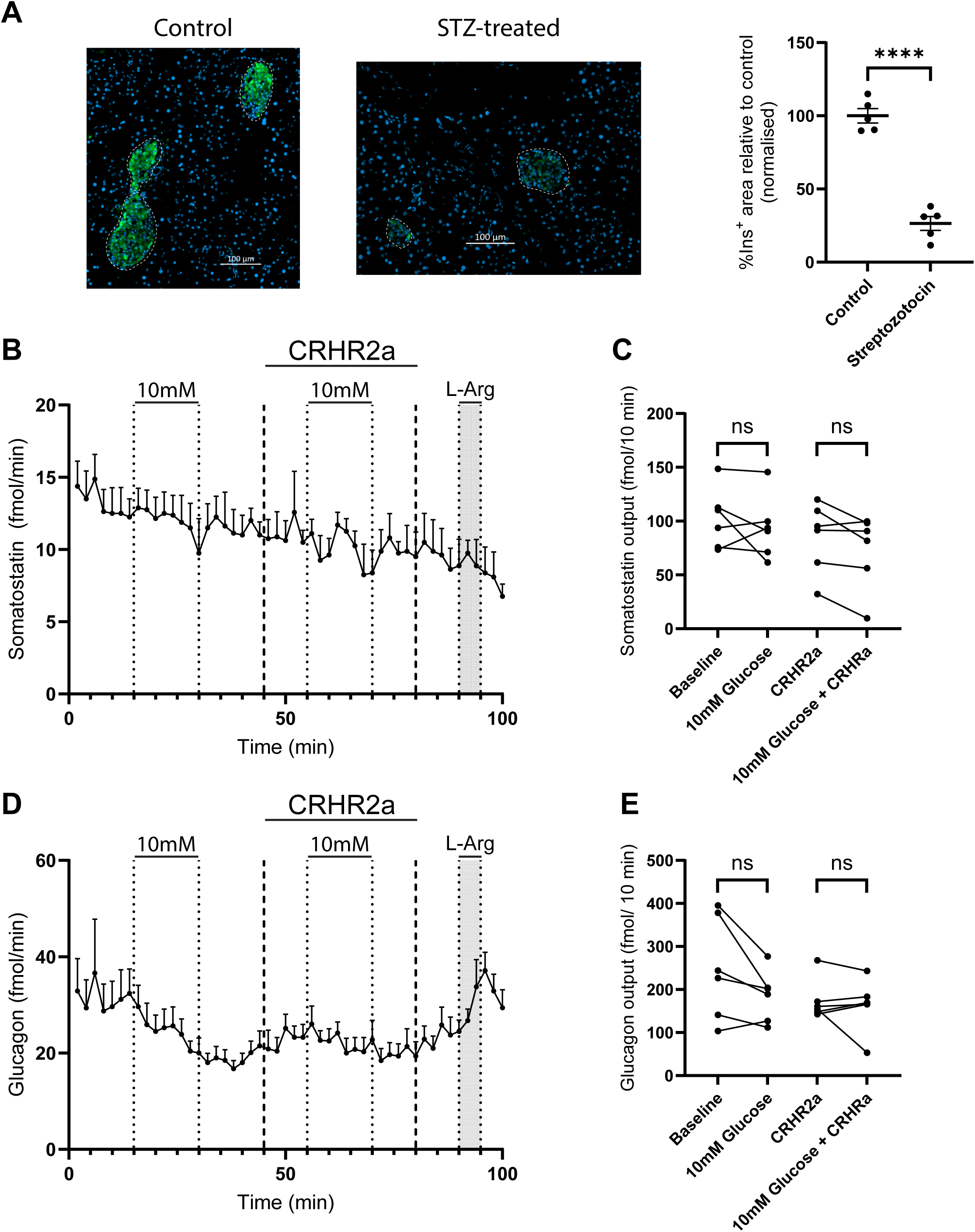
STZ-induced beta-cell ablation impairs glucose-regulated pancreatic hormone secretion. (A) Representative images of a control mouse pancreatic islet and a STZ-treated mouse pancreatic islet immunostained for insulin, together with the quantification of the % of insulin-positive area relative to the control pancreases. Scale bar is 100µm. Data are presented as mean ± SEM, and statistical significance was tested with a Welch’s t-test (**** P<0.0001, N=5 each group). SST (B) and glucagon (D) secretion from perfused pancreases treated with STZ in response to high glucose levels (10mM), with or without the CRHR2 antagonist (K41498, 100nM). SST (C) and glucagon (E) hormone output. The baseline buffer has 3.5mM glucose. Data are presented as mean + SEM in B and D, and as individual points in C and E. N=6. Statistical significance was assessed using a mixed-effects analysis, followed by a Holm-Šídák correction for multiple comparisons. (ns – non-significant).

In STZ-treated mice, SST secretion was not stimulated by 10mM glucose (102.37±11.41 fmol/10min vs 93.75±11.92 fmol/10min, p-value=0.66), and UCN-3 receptor antagonism likewise had no effect (85±13.3 fmol/10min vs 72.75±14.17 fmol/10min, p-value=0.145) (Fig. 5B-C). Glucagon secretion was not significantly reduced at 10mM glucose (248±48.78 fmol/10min vs 185±24.38 fmol/10min, p-value=0.15) and was unchanged during UCN-3 receptor blockage (174.25±19.14 fmol/10min vs 163.37±25.1 fmol/10min, p-value=0.83)(Fig. 5D-E). These findings are compatible with a role for endogenous UCN-3 in glucose-regulated somatostatin secretion, although contributions from other beta-cell-derived signals cannot be excluded, and support that non-beta-cell-derived factors (direct or paracrine) exert an inhibitory tone on glucagon secretion.

## Discussion

In this study, we investigated the role of exogenous and endogenous UCN-3 in the isolated perfused mouse pancreas. We show that UCN-3 strongly stimulates somatostatin and indirectly inhibits glucagon secretion via SSTR activation. We also demonstrate that CRHR2 activation is essential for UCN-3 to exert its effects in the pancreas, as inhibiting CRHR2 prevented both somatostatin stimulation and glucagon inhibition. Importantly, blocking CRHR2 attenuated the acute response of SST and glucagon secretion to hyperglycemia, supporting a physiological role for endogenous UCN-3 signaling in glucose-regulated islet function. Finally, disrupting the beta-cell-derived paracrine environment led to a loss of glucose-stimulated SST secretion, further highlighting the importance of beta-cell factors in regulating delta-cell activity and, consequently, glucagon secretion.

A regulatory role for UCN-3 in the endocrine pancreas has only been considered during the last decade. Initial studies suggested that UCN-3 directly stimulated insulin secretion [22], whereas subsequent work by van der Meulen and colleagues identified UCN-3 as a paracrine activator of pancreatic delta-cells, promoting somatostatin release and thereby inhibiting islet hormone secretion [14]. More recently, our lab demonstrated that UCN-3 inhibits glucagon secretion in the perfused rat pancreas [17], and similar observations were reported in isolated mouse islets [16]. However, the mechanism underlying this glucagonostatic action remained incompletely understood. It was reported that UCN-3’s inhibitory effects on alpha-cell activity were Gαi-dependent, but it was not determined whether this inhibitory pathway was driven by SST. In the present study, pharmacological blockade of somatostatin receptors completely abolished the inhibitory effect of UCN-3 on glucagon secretion, demonstrating that activation of SST signaling is required for this response and supporting a model in which UCN-3 regulates alpha-cell function indirectly by activating delta-cells, leading to subsequent somatostatin release.

Previous studies identified CRHR2 as the cognate receptor for UCN-3 and reported its selective expression in pancreatic delta-cells [14, 23]. Here, we show that in the mouse pancreas, CRHR2 receptor protein is exclusive to delta-cells, consistent with delta cell-exclusive expression. Also, activation of CRHR2 was required for the somatostatinotropic effects of UCN-3, and pharmacological inhibition of CRHR2 increased insulin secretion, supporting a negative feedback loop that controls beta-cell activity[14]. In agreement with these observations, we found that CRHR2 antagonism completely abolished the stimulatory effect of UCN-3 on somatostatin secretion. Furthermore, blockade of CRHR2 prevented UCN-3-induced inhibition of glucagon, demonstrating that CRHR2 signaling is required for both the somatostatinotropic and glucagonostatic actions of UCN-3.

Although the pharmacological actions of exogenous UCN-3 have been previously characterized, less is known about the physiological contribution of endogenous UCN-3 to glucose-regulated islet hormone secretion. In the present study, we show that endogenous UCN-3 contributes to the regulation of SST and glucagon secretion in response to hyperglycemia, as blocking UCN-3 actions blunts the effectiveness of glucose in stimulating SST and inhibiting glucagon secretion. This supports a model in which rising glucose concentration activates a beta-cell-derived pathway that increases delta-cell activity and indirectly inhibits glucagon secretion, thereby resolving the problem that stimulating beta-cell activity with glucose leads to glucagon inhibition, whereas insulin alone does not inhibit glucagon secretion. However, the contribution of UCN-3 to glucagon secretion appears context-dependent. While CRHR2 antagonism substantially reduced the acute effects of glucose on SST and glucagon secretion, it had only a modest effect during sustained hyperglycemia, reducing SST secretion but not affecting glucagon secretion. These data suggest that UCN-3 is particularly important during dynamic changes in glucose levels, whereas other mechanisms may become more important under sustained hyperglycemic conditions. This idea is supported by findings that glucagon secretion is not altered in high-glucose conditions in a constitutive UCN-3 knockout mouse [16], suggesting that compensatory mechanisms maintain appropriate glucagon secretion. Such mechanisms might include direct effects of glucose on alpha-cells, other beta-cell-derived factors, or additional inhibitory paracrine signals. Together, these findings identify endogenous UCN-3 as a physiologically relevant component of the acute glucose-sensing network within the pancreatic islet, whereas the maintenance of glucagon inhibition during sustained hyperglycemia appears to rely on additional regulatory mechanisms.

Somatostatin is a particularly potent inhibitor of alpha-cells [6, 7] and even relatively low concentrations may be sufficient to maintain glucagon suppression. Previous studies have demonstrated that delta-cells remain active at glucose concentrations below the threshold for beta-cell activation [24] and accordingly, somatostatin secretion is detectable even under hypoglycemic conditions [13]. Consistent with this, blockade of SSTR signaling markedly increases glucagon secretion in the perfused pancreas even at extreme hypoglycemic conditions (1.5mM) [25], indicating the existence of a constant tonic somatostatin-mediated inhibitory tone. Although CRHR2 antagonism reduces SST secretion during sustained hyperglycemia, the remaining somatostatin output may still be sufficient to restrain glucagon secretion. These findings suggest that multiple pathways interact to maintain glucagon suppression during prolonged hyperglycemia.

Dysregulated glucagon secretion is a hallmark of both type 1 and type 2 diabetes, yet the mechanisms responsible for this defect remain incompletely understood. It has been reported that islets from patients with type 2 diabetes exhibit decreased UCN-3 expression [14], which may be linked to abnormal glucagon secretion in these patients. To investigate the consequences of disrupting beta-cell-derived paracrine signaling, we examined pancreatic hormone secretion in STZ-treated mice, a model characterized by extensive beta-cell ablation and a type 1 diabetes-like phenotype. Under these conditions, glucose failed to stimulate SST secretion and only weakly inhibited glucagon secretion, indicating that normal delta-cell responsiveness to glucose depends on signals originating from beta-cells. Loss of beta-cell-derived UCN-3 may contribute to this phenotype, as inhibition of the UCN-3/CRHR2 pathway in healthy pancreases attenuated glucose-induced SST secretion. However, beta-cell ablation also disrupts several other signaling pathways that may be involved in regulating delta-cell function. For example, insulin has been proposed to inhibit glucagon secretion and stimulate delta-cell activity through SGLT2-dependent signaling [10], although we have been unable to confirm this [13]. Recent studies have demonstrated direct electrical coupling between beta- and delta-cells through gap junctions, providing another mechanism by which beta-cells could influence delta-cell activity [26, 27]. Consequently, the altered hormone secretion pattern observed in STZ-treated mice is likely the result of disruption of multiple beta-cell-derived pathways rather than solely the loss of UCN-3 signaling. Nevertheless, the ability of CRHR2 antagonism to phenocopy part of this STZ-induced defect in glucose-stimulated SST secretion supports the idea that endogenous UCN-3 constitutes one important component of the beta-cell-to-delta-cell signaling network.

For this study, we used the isolated perfused mouse pancreas, a preparation that preserves the native pancreatic cytoarchitecture, vascularization, and intra-islet communication while allowing dynamic assessment of hormone secretion with high temporal resolution. In addition, the isolated organ preparation permits investigation of pancreatic hormone secretion without interference from extrapancreatic factors. These features represent important strengths of the present study. Nevertheless, several limitations should be acknowledged. First, CRHR2 localization was assessed using an antibody tested in CRHR2-transfected HEK293 cells. Although the staining pattern was consistent with previous reports demonstrating selective Crhr2 protein expression in pancreatic delta-cells, additional validation approaches would strengthen the evidence for the localization of endogenous CRHR2 protein. Second, the role of the UCN-3/CRHR2 pathway was examined using pharmacological antagonists rather than genetic loss-of-function models. Finally, interpretation of the STZ experiments is limited by the fact that beta-cell ablation disrupts multiple beta-cell-derived signals in addition to UCN-3. Therefore, the phenotype observed in STZ-treated mice cannot be attributed exclusively to loss of UCN-3 signaling.

In summary, our results support a model in which beta-cell-derived UCN-3 acts via CRHR2 to stimulate delta-cell SST secretion, thereby contributing to the suppression of glucagon secretion. We further demonstrate that endogenous UCN-3 participates in the acute islet response to hyperglycemia, identifying the UCN-3/CRHR2 pathway as a physiologically relevant component of intra-islet communication. While this pathway contributes to the rapid adaptation of glucagon secretion to changes in glucose concentration, additional mechanisms seem to contribute to glucagon suppression during sustained hyperglycemia. Finally, the impaired hormone responses observed following beta-cell ablation underscore the importance of beta-cell-to-delta-cell signaling in regulating islet function and suggest that loss of UCN-3 may contribute to the dysregulation of glucagon secretion observed in diabetes.

## Data availability

The datasets generated and/or analyzed during the current study are available from the corresponding author upon reasonable request.

## Acknowledgements

The authors thank Søren Petersen (Department of Biomedical Sciences, University of Copenhagen) for technical assistance with the cell transfection experiments. We also thank BMI Histolab for tissue processing and sectioning of pancreatic samples for histology. Finally, we acknowledge the Core Facility for Integrated Bioimaging, Faculty of Health and Medical Sciences, University of Copenhagen, for access to the microscope.

## Disclosure of AI use

Claude Sonnet 5.5 (Anthropic) was used to assist in writing R code for data analysis. In addition to Claude, Microsoft Copilot (GPT-5 chat model) was used to assist with the text’s language, readability, and proofreading. The authors reviewed all the AI-generated output and take full responsibility for the content of the manuscript.

## Author contribution

JVU, DBA, and JJH designed research. JVU and DBA performed research. JVU, DBA, GDG, TM, MMR, and JJH analyzed and interpreted the data. JVU, DBA, and JJH wrote the manuscript. All authors read, edited, and approved the final manuscript.

## Authors’ relationships and activities

The authors declare there are no conflicts of interest associated with this manuscript.

## Abbreviations

AUC: Area Under the Curve
BSA: Bovine Serum Albumin
cAMP: Cyclic Adenosine Monophosphate
CRHR2: Corticotropin-Releasing Hormone Receptor 2
CRHR2a: CRHR2 Antagonist
EtOH: Ethanol
FBS: Fetal Bovine Serum
GABA: Gamma-Aminobutyric Acid
HEK293: Human Embryonic Kidney 293 Cells
i.p: Intraperitoneal
L-Arg: L-Arginine
mCRHR2: Mouse CRHR2
PFA: Paraformaldehyde
SEM: Standard Error of the Mean
SGLT2: Sodium-Glucose Cotransporter 2
SST: Somatostatin
SSTR: Somatostatin Receptor
SSTRa: Somatostatin Receptor Antagonist Cocktail
STZ: Streptozotocin
UCN-3: Urocortin-3

---

**Supplemental Figure 1.**
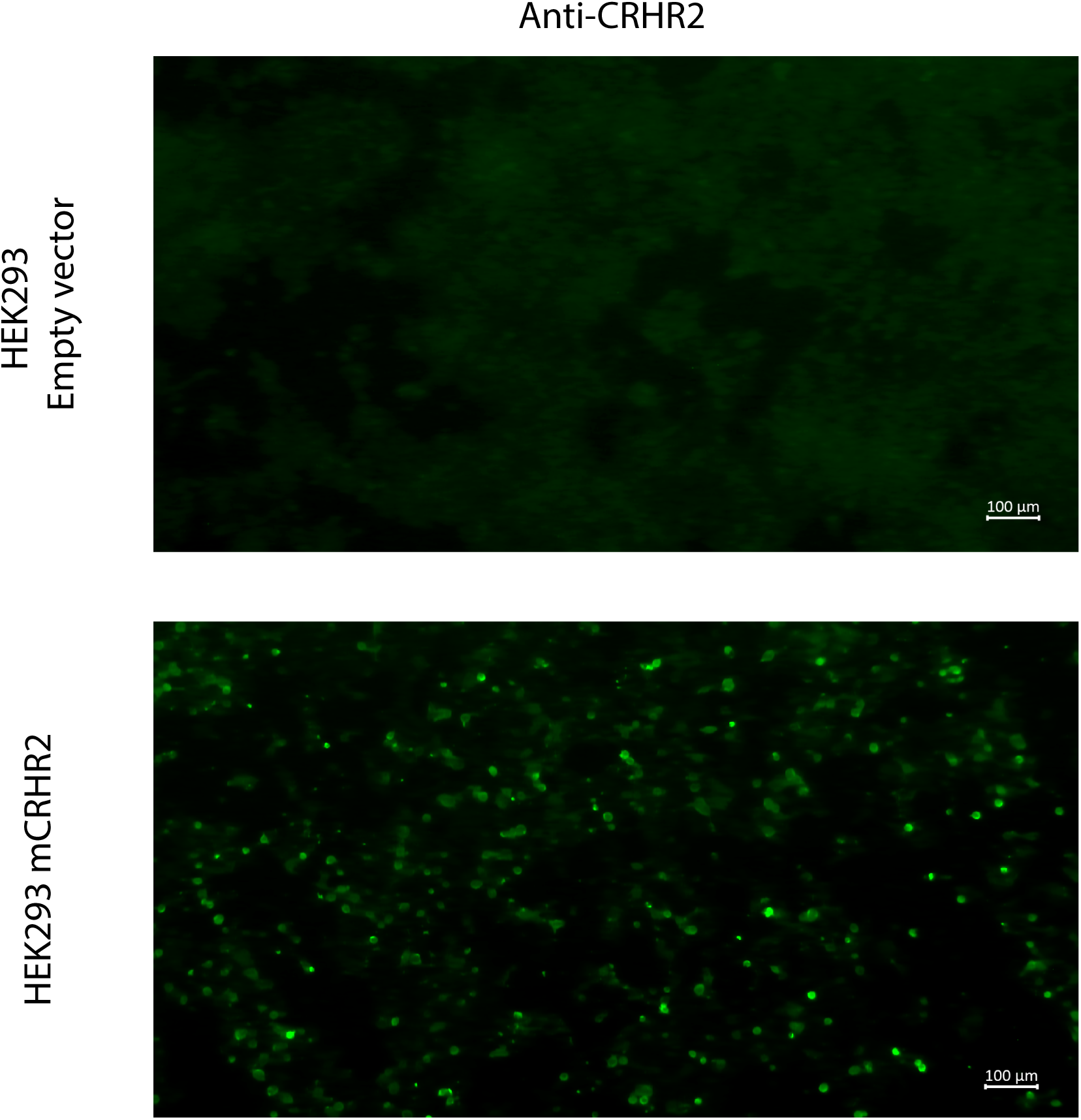
Representative immunofluorescence images of HEK293 cells transfected with empty vector (upper panel) or with mouse CRHR2 (mCRHR2) (lower panel) and stained with an anti-CRHR2 antibody. Scale bars: 100μm.

|  | Antibody Target | Technical information | Dilution | Host | Tissue preparation |
|---|---|---|---|---|---|
| Primary | Insulin | In-house antibody.<br>Ab code: 2006-4 | 1:1000 | Guinea-Pig | Paraffin |
|  | Somatostatin | Santa-Cruz Biotechnology,<br>G-10, monoclonal | 1:100 | Mouse | Paraffin |
|  | CRHR2 | Abcam, ab150510,<br>polyclonal | 1:100 | Rabbit | Paraffin |
| Amp. | Amplifier antibody | Vector laboratories, DK-1488,<br>Anti-rabbit | RTU* | Goat | Paraffin |
| Secondary | Alexa fluor 488<br>Anti-Guinea-Pig | ThermoFisher Sientific,<br>A11073 | 1:200 | Goat | Paraffin |
|  | Alexa fluor 568<br>Anti-Mouse | ThermoFisher Sientific,<br>A10037, polyclonal | 1:200 | Donkey | Paraffin |
|  | Vecta fluor 488 | Vector laboratories, DK-1488,<br>Anti-goat | RTU* | Goat | Paraffin |

\*RTU: Ready to use

